# The platyrrhine primate *Cebus imitator* uses gaze to adjust grasp posture for food handling and withdraw to the mouth

**DOI:** 10.1101/2023.06.22.546193

**Authors:** Ian Q Whishaw, Megan Mah, Julia G. Casorso, Evin Murillo Chacon, Janine Chalk-Wilayto, Myra Laird, Amanda Melin

## Abstract

Orienting a food item held in the hand to withdraw it to the mouth for eating is mediated by vision in catarrhine anthropoids and by nonvisual strategies in strepsirrhines. The present study asks whether vision contributes to the withdraw in a platyrrhine anthropoid, a member of a monophyletic primate suborder whose stem group diverged from catarrhines about 40 million years ago. We examined gaze and hand use in arboreal fruit picking by the Costa Rican capuchin, *Cebus imitator*, a platyrrhine known for its skilled object-handling and tool use. Videos clips of reach, grasp and withdraw movements and associated gaze were examined frame-by-frame to assess hand shaping and sensory control of eating. *Cebus imitator* used vision and touch to reach for and grasp food items with precision or whole hand grasps. They used vision to orient food items held inhand into a precision grip and their withdraw of food items to the mouth was assisted with a vertically oriented hand. The conjoint use of vision, grasp and hand posture by capuchins is discussed in relation to the staged evolution of visual manipulation of objects, one of which is featured in this platyrrhine primate.

## Introduction

The evolutionary origins of the visual control of reaching for eating, tool use, and other skilled hand activities by humans is unresolved. In addition, contemporary visuomotor channel theory proposes that skilled forelimb use is composed of a number of sub movements; for example, for eating, the reach, the grasp, and the withdraw of a food item to the mouth (Jeannerod,1981; Jeannerod et al., 1995, 1998; Grant & Conway, 2019; Sartori et al, 2015; Whishaw et al, 2016). This organization implies that each channel and subcomponent movement has its own evolutionary history (Karl and Whishaw, 2013). Insight into the visual control of hand use related to the sub movements of reaching can be obtained by examining these movement in the many animal species that use their hands for food manipulation and especially in extant primates that use vison for food manipulation (Iwaniuk et al., 1998; Sustaita et al, 2013; Whishaw and Karl, 2014; 2019). Catarrhine primates - as represented by macaques (*Macaca*), anthropoid members of the subfamily Cercopithecinae - pick up food items using hand and finger shaping strategies, including the pincer and precision grasps, that are appropriate to the size and shape of food items (Bishop 1964; Christel 1993; Christel and Fragaszy 2000; Macfarlane and Graziano 2009; Marzke *et al*. 2015; Pouydebat *et al*. 2008; Scott 2019). They also visually examine food objects that they hold in the hand and use vision to orient food items to place them in the mouth, as do other catarrhine anthropoids including humans (Hirsche et al, 2022). The co-occurrence of visually controlled precision grasping and food manipulation inhand for mouth placement in catarrhines suggests that the visual control of these behaviors may be related. Comparative studies support this conclusion as nonprimate species do not use visually mediated hand shaping for reaching or for oromanual behavior including handling food items for placement in the mouth. Nonprimate species pick up a food item by mouth and/or reach with the mouth to take a food item held in the hand (Ivanco et al., 1996; Iwaniuk et al., 1998; Sustaita et al, 2013; Whishaw & Coles, 1996; Whishaw et al., 1998, 2018, 2020). Strepsirrhines, an early branching suborder of primates, also do not make visually mediated hand shaping movements for picking up food or for withdraw movements (Reghem *et al*. 2011; 2023; Perrenoud *et al*. 2020; Pouydebat et al., 2008; 2009). Some strepsirrhine species may visualize and pick up smaller food items by mouth (Peckre et al, 2016) and all species use their mouth to take food from the hand and use nonvisual strategies for hand to mouth food transfer (Peckre et al, 2023).

The use of vision for object grasping and for inhand withdraw movements to place food into the mouth in catarrhines, vs. the absence of similar visual use in strepsirrhines, raises questions related to the evolutionary origins of the visual contribution to these movements. Did visual control of the grasp and the withdraw coevolve, did visual control of the grasp contribute to the subsequent visual control of the withdraw, or was this sequence of events reversed? Evidence related to these questions could be obtained by examining the sensory control of the grasp and the inhand withdraw in the other major radiation of anthropoids, the monkeys of the Americas (Platyrrhini). The stem group of this monophyletic primate suborder split from catarrhines about 40 million years ago (Arnold et al, 2010; Kay et al, 1997; Kissling et al, 2015), and so the strategies that they use in feeding could provide insights into the evolution of visually mediated reaching and withdraw movements that are absent in strepsirrhines but present in catarrhines. Additionally, examination of the feeding behavior of platyrrhines might shed light on whether the acquisition of these behaviors was a singular event or whether one preceded the other. Thus, the objective of the present study is to determine whether a platyrrhine primate uses vision to orient food items held in the hand for placement in the mouth. The answer to this question is relevant to understanding the use of vision in object control by the hand more generally.

We studied a population of white-faced capuchins, *Cebus imitator*, from Costa Rica, members of the subfamily Cebinae. Amongst platyrrhines, Cebinae are described as especially skillful in hand use (Christel & Fragazy, 2000; Costello & Fragazy, 1988; Melin et al, 2022; Spinozzi et al, 2003; Truppa et al, 2019a,b; 2020). They are also representative of a species foraging in a termineal branch niche, a niche proposed to be central to the evolution of the visual control of hand (Cartmill, 1972, 2012; Sussman & Raven, 1978; Sussman et al., 2013; Scott, 2019). Wild, unprovisioned *Cebus imitator* were filmed as they foraged in trees for a variety of fruits. The videos were trimmed to include examples of fruit picking and eating and were then examined frame-by-frame to assess gaze in relation to the reach, grasp and the handling/withdraw movements of bringing food items to the mouth using previously described methods (Hirsche et al, 2022; Peckre et al, 2023).

## Materials and methods

### Study populatiion

The feeding behavior of *Cebus imitator* was filmed in the Sector Santa Rosa (SSR), Área de Conservación Guanacaste (ACG) in northwestern Costa Rica (10°450–11°000 N, 85°300– 85°450 W). The study population consisted of 16 animals, 5 adult males, 3 adult females, 2 subadult males, 3 juvenile females, and 3 juvenile males. Filming consisted of short (<1–10 min) continuous video samples following a published protocol [22] with strict out-of-site rules, such that recording of behavior was done when there was a relatively unobstructed view of the focal monkey’s feeding behavior. Individuals were sampled opportunistically, based on visibility, but observations rotated among sex and age classes to sample evenly across the population. This research adhered to the laws of Costa Rica, the United States, and Canada and complied with protocols approved by the ACG (R-SINAC-ACG-PI-027-18) (R-025-2014-OT-CONAGEBIO), by the Canada Research Council for Animal Care through the University of Calgary’s Life and Environmental Care Committee (AC19-0167), and by Mercer University’s Institutional Animal Care and Use Committee (A2002003).

### Video Recording

Video recording at 30 frames per sec (fps) provided *ad libitum* samples of eating behavior of the *Cebus*. Video-recorded data were collected using Lumix DC-G9, Sony model FDR-AX53, and Olympus OM-D E-M1 camcorders.

### Food items

Fruit items eaten by the capcuhins included *Bromelia pinguin* (wild pineapple, Piñuela; round, 3-4 cm), *Spondias mombin* (hog plum, Jocote jobo; ovoid, 3.5-5 cm long, 2.5-3 cm wide), *Bursera simaruba* (gumbo-limbo, Indio desnudo; round, 1-1.5 cm), *Trichilia martiana (*Manteco; round, 1-1.5 cm), *Genipa americana (*Guaitil; round, 6-8 cm), *Diospyros salicifolia* (persimmons, Lorito; round, 1.5-2 cm), *Byrsonima crassifolia* (nance; round, 1-1.5 cm), *Luehea candida (*Guácimo molenillo; ovoid, 6-8 cm long, 3.5-5 cm wide), *Ficus cotinifolia (fig*, Higuerón; round, 1-1.5 cm), *Ficus ovalis (fig*, Higuerón; round, 1-1.5 cm), *Ficus hondurensis* (fig, Higuerón; round, 1-1.5 cm), and *Stemmadenia obovata (*Huevos de burro; ovoid, 6-7 cm long, 3-4 cm wide).

## Behavioral analysis

### Video analysis

The video recordings of *capuchins* eating were examined frame-by-frame using *Quicktime 7.7 (*https://support.apple.com/en-ca/guide/quicktime-player/welcome/mac) on an Apple computer. IQW performed the behavioral scoring detailed below based on previously described methods (Hirsche et al, 2022; Peckre et al, 2023). The scoring systems have been performed by frame-by-frame video analyses using this scoring system have yielded an inter-scorer reliability coefficient of 0.96 (Hallgren, 2012). Every feeding sequence, in which a capuchin reaches for, grasps, withdraws and places a food item in the mouth was analyzed. Because animals were reaching through leaves and adjusting posture, some component movements of a reach were visible and others were obscured, nevertheless, the observable components were always scored.

### Behavioral analysis

1. *Body posture.* Body posture associated with reaching movements was scored on a 5-point scale (Peckre et al, 2023; Laird et al, 2022):

> 0 – A score of “0” was given if the long axis of an animal’s body was in a horizontal orientation relative to the food target for which it was reaching.
>
> 1 – A score of “1” was given if the torso was in about a 45^0^ upright orientation from the horizontal.
>
> 2 - A score of “2” was given if the long axis of an animal’s body was in a vertical upward orientation relative to the food target for which it was reaching.
>
> -1 – A score of “-1” was given if the torso was in about a 45^0^ downward orientation from the horizontal.
>
> -2 - A score of “-2” was given if the long axis of an animal’s body was in a vertical downward orientation relative to the food target for which it was reaching.
2. *Stance*. Stance was scored on a 5-point scale in relation to the number of limbs that were supporting the body during a reach for food and withdraw of the food (Reghem et al., 2011; Whishaw et al., 1988). Although the tail contributs to support, it was not included in the stance score:

> 0 – designated that an animal was draped over a branch or suspended by the tail.
>
> 1 – designated that an animal was standing on one limb.
>
> 2 – designated that an animal was supported on two limbs, usually in a sit posture.
>
> 3 – designated that an animal was supported on three limbs, usually reaching with the fourth limb.
>
> 4 – designated that an animal was supported on four limbs and was thus reaching with its mouth.
3. *Head orientation*. The extent to which the head contributed to the withdraw of food by reaching for a food item was rated on a 5-point scale (Hirsche et al, 2022; Peckre et al, 20023):

> 0 – the head was advanced to the food and the food was grasped with the mouth
>
> 1 – the nose was placed near the target as the item was grasped by hand and brought to the mouth
>
> 2 – the hand and the mouth were brought toward each other such that the withdraw movement was accomplished equally by the hand and mouth.
>
> 3 – most of the withdraw was accomplished with the hand with only a small orienting movement made by the head toward the hand.
>
> 4 – the head was not advanced toward the food or withdrew as the hand brought the food toward the mouth.
4. *Hand use for reaching.* Two types of hand use for reaching were documented by counts of occurrences (Fragaszy, 1968; Spinozzi et al, 2004; Truppa et al, 2019):

1. *Single hand use*: single hand use involved a capuchin advancing only one hand to grasp an item with the other hand used for either supporting weight or grasping a branch for balance.
2. *Bilateral hand use*: Bilateral hand use involved a capuchin using one hand to grasp a branch containing a fruit item and manipulating the branch into a position from which the other hand could grasp the fruit on the branch.
5. *Mouth grasping from the hand.* The way in which the mouth grasped food items from the hand was documented with counts of occurrences (Hirsche et al, 2022):

1. *Incisor grasp*: incisor grasp consisted of the mouth opening and grasping a food item with a precise grasp using the incisor teeth, usually with the food item presented to the front of the mouth.
2. *Premolar grasp*: premolar grasps consisted of the food item being grasped by the premolar teeth, usually with the food item presented to the side of the mouth.
6. *Adjunct oral movements.* Associated with reaching behavior, the capuchins made two kinds of oral movements during reaching and these were documented in animals eating *Ficus ovalis*, a grape sized fig (Vainio et al, 2019):

1. *Gapes*: gapes were mouth openings in which neither the tongue nor teeth other than the canines were visible
2. *Spits*: spits involved an animal opening its mouth and spitting out food
7. *Reaching movements.* A reaching movement consisted of advancing a hand to a food item, grasping the item, and withdrawing the food item to the mouth so that it could be grasped by the mouth. The component movements of a reaching movement were scored as follows (Karl and Whishaw, 2013):

1. *Reach.* A reach consisted of an advance of the hand to a food item, and its duration was measured by counting video frames that started with the first movement of advancing the hand toward the food and ended with the frame on which the advancing movement ended.
2. *Grasp.* A grasp consisted of the movement closing the digits so that the food item was held in the hand, and each grasp was measured by counting frames, beginning with the last frame of the reach to the first frame where hand completed the grasp and moved toward the mouth.
3. *Immediate Withdraw.* An immediate withdraw was a movement that brought a food item to the mouth without a pause, and was measured by counting video frames, beginning with the frame where the grasp ended to the frame at which the hand stopped to transfer the food item to the mouth.
4. *One-handed food holding*. A one-handed food holding movement consisted of holding a food item in one hand before bringing the food item to the mouth.
5. *Two-handed food holding*. A two-handed food holding movement consisted of holding a food item with both hands before bringing it to the mouth.
6. *Inhand withdraw*. An inhand withdraw consisted of bringing a food item that been held in one or both hands to the mouth. The duration of an inhand withdraw was measured by counting frames, beginning with the first movement of the hand holding a food item toward the mouth until the hand came to a stop to place the food in the mouth.
8. *Hand grasps.* The capuchins made different types of hand grasps when initially grasping and handling food items (for descriptions of capuchin hand grasps see Felix et al, 2015; Fragaszy, 1968). Hand posture was recorded both when a food item was picked up and when a food item was transferred to the mouth. Grasps were divided into two categories:

1. *Precision grasp*: a food item was grasped or held mainly between the first two fingers (pollex and index finger) and might also be pressed against the palm.
2. *Whole hand grasp*: a food item was grasped or held between the pollex and two or more of the other fingers and also pressed against the palm.
9. *Gaze.* When food items were grasped, if the head of an animal was oriented toward the food at the time that the grasp occurred, that was taken as a sign that the animal was looking at the food; item i.e., visually engaged or gaze anchoring as defined by Posner et al. (1987). The following head/eye orienting behaviors were quantified:

1. *Engage*: a movement that resulted in the face/eyes being directed toward the food item before or as it was grasped. Gaze duration was measured with frame counts from the point that the head/eyes were directed to a food item to the frame on which they began to move away from that orientation, or the capuchin blinked.
2. *Disengage*: a movement that resulted in the face/eyes being directed away from the food item that had been grasped (cf. de Bruin et al, 2008).
3. *Eye blink*: Eye blinks were rapid closing and opening of the eyelid. The occurrence of a blink was noted for those video recordings for which a view of the eyes was adequate and as a proportion of all withdraw movements (cf. de Bruin et al., 2008).

### Statistical analyses

The effect of withdraw movements, duration, and visual attention according to the food item size, subject age, and subject sex was analyzed using Generalizes Linear Model ANOVAs with repeated measures and t-tests for paired samples using the program SPSS (v.24.0.0). Mauchly’s tests of sphericity were used to compare variance and when the assumption of sphericity was not met, analysis was made with Huynh-Feldt correction. Results are reported as mean ± standard error. A *p* value of < 0.05 was considered statistically significant and a partial eta squared (η*P*^2^), a descriptive measure of the strength of association between the independent and dependent variables, was used to measure the effect size (Kelley & Preacher, 2012). Counts of behaviors including posture, grasping type, engage and disengage, blinking and eye direction are reported as the percent of the number of observations that were made. The relationship between the duration of the ground-withdraw and inhand-withdraw movements to head-engage and disengage duration were assessed using the Pearson product-moment correlation and compared with Chi-square tests.

## Results

The results were obtained from 390 video clips each ranging from 15 sec to 10 min, and which together comprised 9.44 hr of video. Frame-by-frame inspection of the video provided 598 instances of reaching, a movement sequence comprising a reach by hand for a food item, a grasp of the food item, and the withdraw of the food item to the mouth for eating. Figure 1 illustrates that the capuchins were adept in moving amongst terminal branches of trees and supporting themselves with various limb and prehensile tail configurations, all the while reaching, grasping, and withdrawing fruit items to the mouth for eating. Feeding principally occurred in the terminal branches of trees, a location described as a fine-branch niche (Cartmill, 1972, 2012; Sussman & Raven, 1978; Sussman et al., 2013; Scott, 2019). A few instances of animals coming briefly to the ground to pick up food items were observed, but the animals quickly returned to a tree to eat, consistent with the aboreal proclivities of this species (Fragazey et al, 2004). When a part of the animal was hidden from view by branches or leaves, those parts of the reaching act that were visible are nevertheless described. Consequently, the following descriptions include a different number of observations depending upon the component movement observed. Some of the animals’ feeding movements differed depending upon the fruit and size of items it was eating, and these examples are described as a subset of reaching observations. Given that the data are obtained from the spontaneous behavior of the animals in trees, the movements of reaching were associated with a variety of body movements. Body movements were related not only to the movement of the animals themselves but also to the branch movements produced by the animal’s postural changes, the wind, and movements of other animals. Because the kinematics of reaching were not being measured, behavioral observations could be scored despite the complexity of substrate/body movements. The measures of reaching described below all involved reaching acts for obtaining food and bringing it to the mouth for eating. Reaching acts associated with bringing a food item to the nose for sniffing, grasping a food item and not picking it, and grasping objects that were released or dropped were not documented.

**Figure 1.**
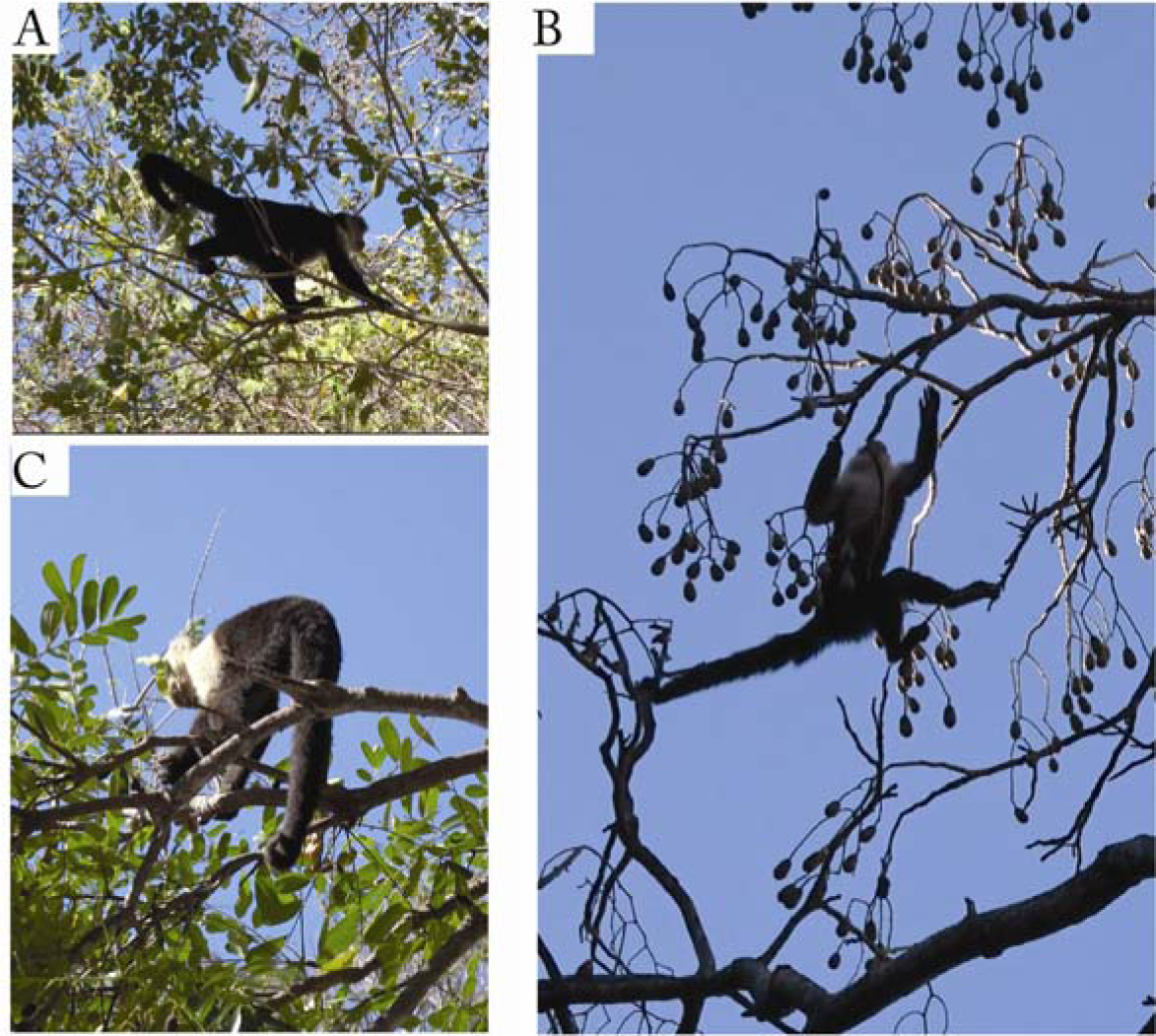
Foraging habitat. Capuchins foraging in: (A). *Ficus cotinifolia*; (B). *Cedrela odorata* (C). *Simarouba glauca*). Note that foraging involved moving amongst relatively small distal branches of trees, in what is termed a terminal branch niche.

### Favored reaching posture, stance, and head orientation

Figure 2 summarizes the body posture, stance, and head orientations observed in 598 reaching observations. Figure 2A illustrates the most frequently occurring postural configurations associated with *Cebus imitator* reaching. Posture features the long axis of the body in an oblique orientation, stance is sitting on the haunches supported by the hind limbs, and the withdraw is made by a single hand bringing a food item to the mouth. The numerical notation used to define posture is depicted by the numbers to the right of the cartoon capuchin (Pessina et al, 2019). Figure 2B illustrates the most frequent posture had the long axis of the torso noted by orientation “1” in Figure 2A. Less frequently used postures included the body in a horizontal position and in a vertical position with the head up or down.

**Figure 2.**
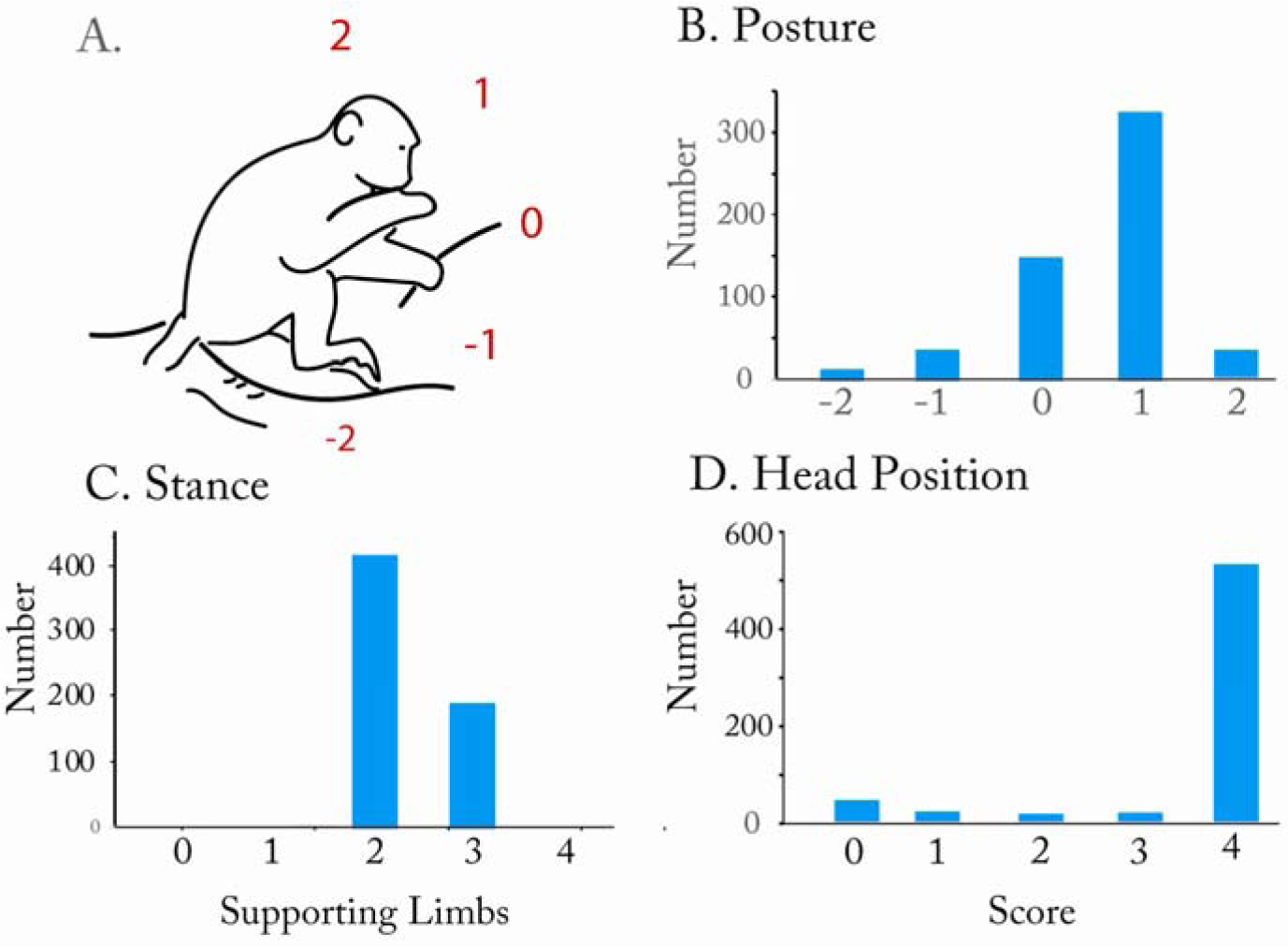
Postural features of capuchins when consuming fruit. (A). An illustration summarizes average postural features with an animal: sitting on its haunches with a trunk angle of approximately 45 degrees, supporting itself on two hind limbs, bringing a food item to the mouth with one hand, with the head upright and drawing away from a hand bringing food to the mouth (numbers refer to body angle, 0=horizontal, 1=angle of 45 degrees, 2+upright angle of 180 degrees, -1 angle of -45 degrees, -2 downward angle of 180 degrees). (B). Posture, relative number of times different eating and reaching postures were used. (C). Number of supporting limbs used by animals when eating and reaching. (D). The number of times that different head configurations were used to bring food to the mouth, with a score of “0” indicating that a food item was reached for with the mouth and a score of “4” indicating that the hand brought a food item to the mouth where it was taken with a discrete bite.

Figure 2C illustrates that the most frequently occurring stance was sitting on the haunches (see Laird et al, 2022 for a description of the relationship between substrate and food type in capuchins), but less frequently animals could be standing on four limbs, three limbs, two limbs and even one limb. A sitting posture is described as an euarchontoglire trait that has been documented in animals feeding on a horizontal surface (Reghem et al., 2011; Whishaw et al., 1988). Figure 2D illustrates that when a food item was brought to the mouth by a hand, the head moved upward and did not advance to take the food from the hand. Thus, the only movement of getting the food to the mouth involved the hand. Instances when the head was directed to an item did occur, the most frequent of which had an animal holding a stick with both hands and reaching with its mouth to chew, or when an animal was attempting to break a food item loose from its stem when grasping it by the mouth, and these behaviors were not counted or analyzed further.

### Variability in reaching and reach component duration

Figure 3A illustrates the distribution of reaching durations (the combined reach, grasp and immediate withdraw). A total of 88 reaching acts were measured from adult animals picking *Ficus ovalis* (a grape sized fig). All of the reaching acts that were measured were those associated with an immediate withdraw; i.e., the food item is brought directly to the mouth after grasping. The vertical dotted lines in Figure 4A give the average durations of the subcomponent movements of reaching: the reach, the grasp, and the withdraw. As Figure 3A illustrates, reaching duration was variable, lasting from less than 1 sec to more than 3 sec, and variability was featured in each of the component movements. Factors influencing durations likely included the distance that an animal was reaching, the involvement of the nonreaching hand in manipulating a branch containing a target item into a location from which it could be grasped by the reaching hand, the movements of the hand over and around vegetation and a target item in order to obtain purchase, and even the slowing of a movement to achieve adjunct movement such as the spitting out of food to make way for a new food item to be placed in the mouth (see below).

**Figure 3.**
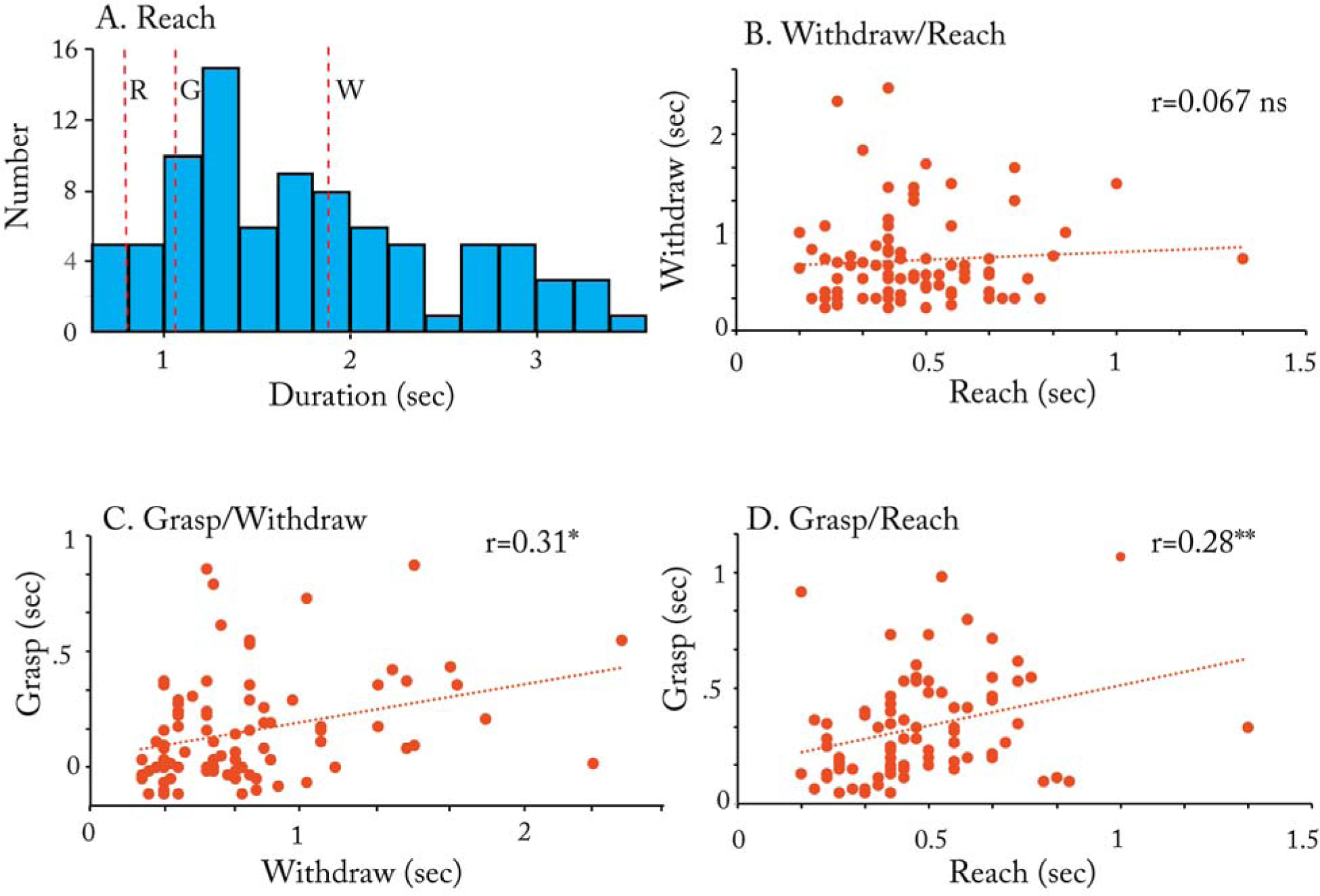
Reach and reach component time. (A). Distribution of all total reach times, with vertical dotted lines indicating the mean duration of the reaching components: reach (R), grasp (G) and withdraw (W). (B). Relationship between mean withdraw duration and mean reach duration. (C). Relationship between mean grasp duration and mean withdraw duration. (D). Relationship between mean grasp duration and mean reach duration. Note: Large variation in reaching times and component times associated with low correlation value suggest independence of the component movements of reaching.

**Figure 4.**
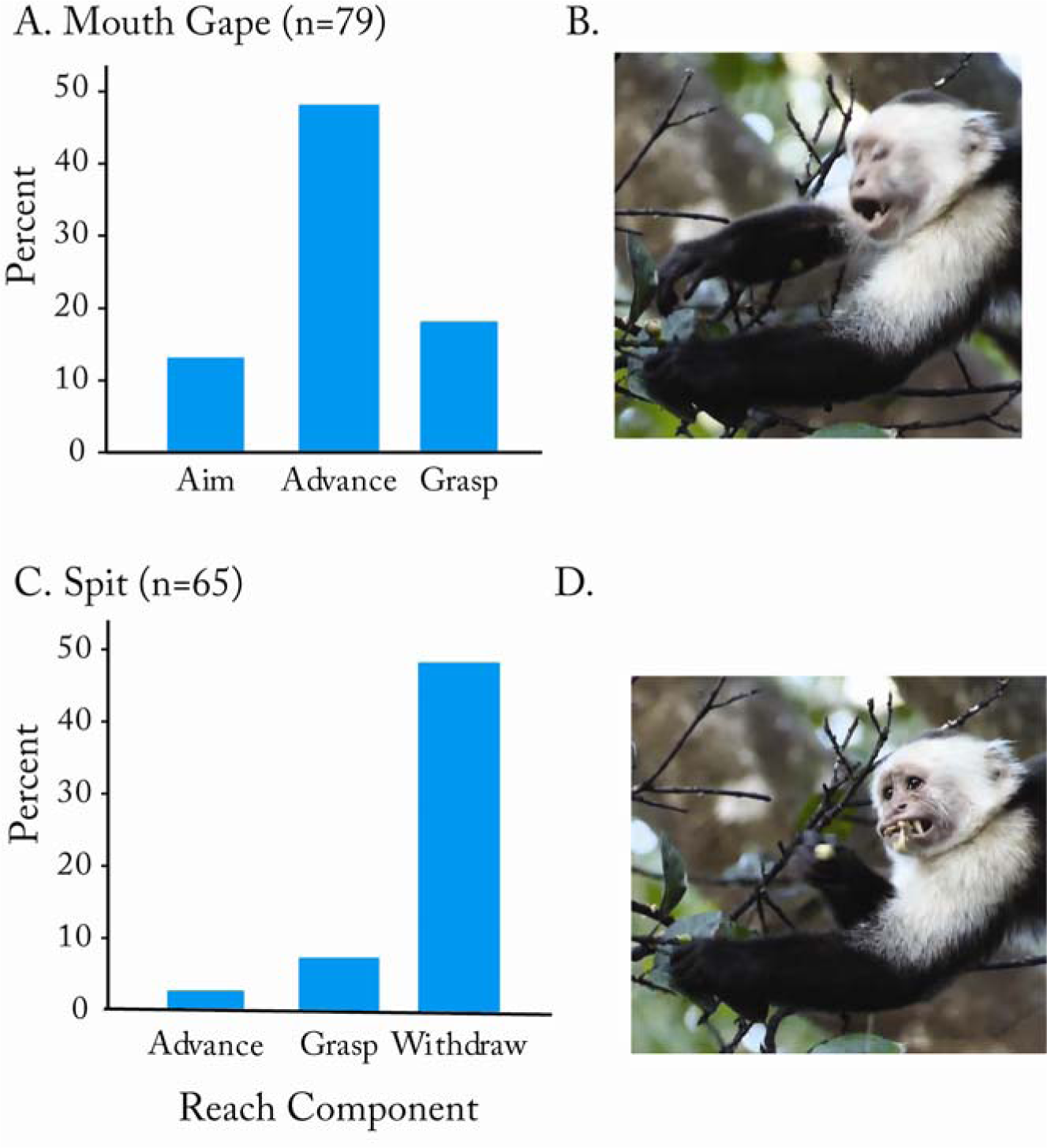
Adjunct oral movement associated with reaching for *Ficus ovalis. (*A). Percent of observations during which mouth gapes occurred during different component movements of the reach as the hand advances to the food. (B). A capuchin named Badger (BD) makes a mouth gape that occurred as the hand was advanced to grab a fruit item. (C). Percent of observations during which spitting movements of the mouth occurred with different components of an immediate withdraw of a fruit item to the mouth. (D). A capuchin named Badger (BD) makes a spitting movement that occurred as the hand was withdrawn to the mouth.

We examined correlations between the reach, grasp, and withdraw (Figures 3B-D). The correlation between the durations of the reach and the withdraw was not significant, r(86)=0.067, p>0.05. Although there were significant correlations between the grasp and the reach durations, r(86)=0.31, p<0.05, and the grasp and the withdraw, r(86)=0.28, p<0.05, inspection of the figures confirm that these correlations are not high. The correlations suggest that, although the reach, grasp and withdraw always occur in the same order, the duration of each movement is variable and is associated with factors other than those related to the duration of the other movement components. Taken together, the variability in movement component durations show that there is substantial autonomy and flexibility in each of the component movements of capuchin reaching.

### Adjunct oral movement accompany reach component movement

Associated with both reaching for a food item and withdrawing a food item to the mouth, the capuchins were observed to make oral movements (for a review of many reports of adjunct oral movements accompanying hand movements, see Vainio, 2019). These movements were documented as capuchins foraged for *Ficus ovalis* and are illustrated in Video 1. As shown in Figure 4A, for 79% of 98 reaches, the capuchins made a mouth gape as illustrated in Figure 4B. The majority of mouth gapes occurred as the hand was advanced to the fruit (48%). Less frequently, mouth gapes occurred as the hand was in the aim position, raised but not yet advancing (14%), or as the hand was positioning to grasp (18%). Mouth gapes did not occur on the withdraw.

As illustrated in Figure 4C, spitting food out of the mouth was observed for 57% of 98 withdraw movements, and spitting occurred usually as the hand approached the mouth (Video 1). During spitting, the hand often visibly slowed its approach to the mouth, apparently so that the animal could clear its mouth to accept a new fruit item as shown in Figure 4D. Food spitting behavior was mainly associated with eating *Ficus ovalis*, suggesting that there were some portions of the fig that were being rejected.

### Gaze and touch strategies are used for the grasp

The contribution of gaze to the grasp of a fruit item was assessed by counting the total frames (from the time the wrist stopped its forward motion toward the target to the time the wrist began a reverse movement to bring the hand to the mouth) and the associated number of frames during which the head was oriented to the target with gaze directed to the target. Figure 5A illustrates the relation between gaze duration and total time taken to grasp a fruit item. There was a significant relationship between gaze duration and grasp time, r(107)=0.50, p<0.001, but both gaze and grasp duration were variable, lasting from less than half a second to as long as one second. Nevertheless, as is illustrated in Figure 5b, about 50% of grasps were not associated with gaze being directed at the food object during grasping and about 15% of grasps were associated with touches that contributed to orienting the hand to grasp (see Video 2). Thus, for about half of all grasps, gaze was not directed to the food item as it was grasped. This may be why many grasps were associated with hand orienting movements seemingly mediated by touch (for a description of the use of touch in foraging capuchins, see Melin et al, 2022). Figure 5C and Figure 5D show examples of touch-associated grasps. Figure 5C shows a photo from a video sequence in which a capuchin contacts a fig with the second finger and maintains contact as it slides its finger over the object before grasping. Figure 5D shows a photo from a video sequence in which a capuchin moves its palm across a fig as if locating the fig, and then reverses its trajectory to grasp the fig.

**Figure 5.**
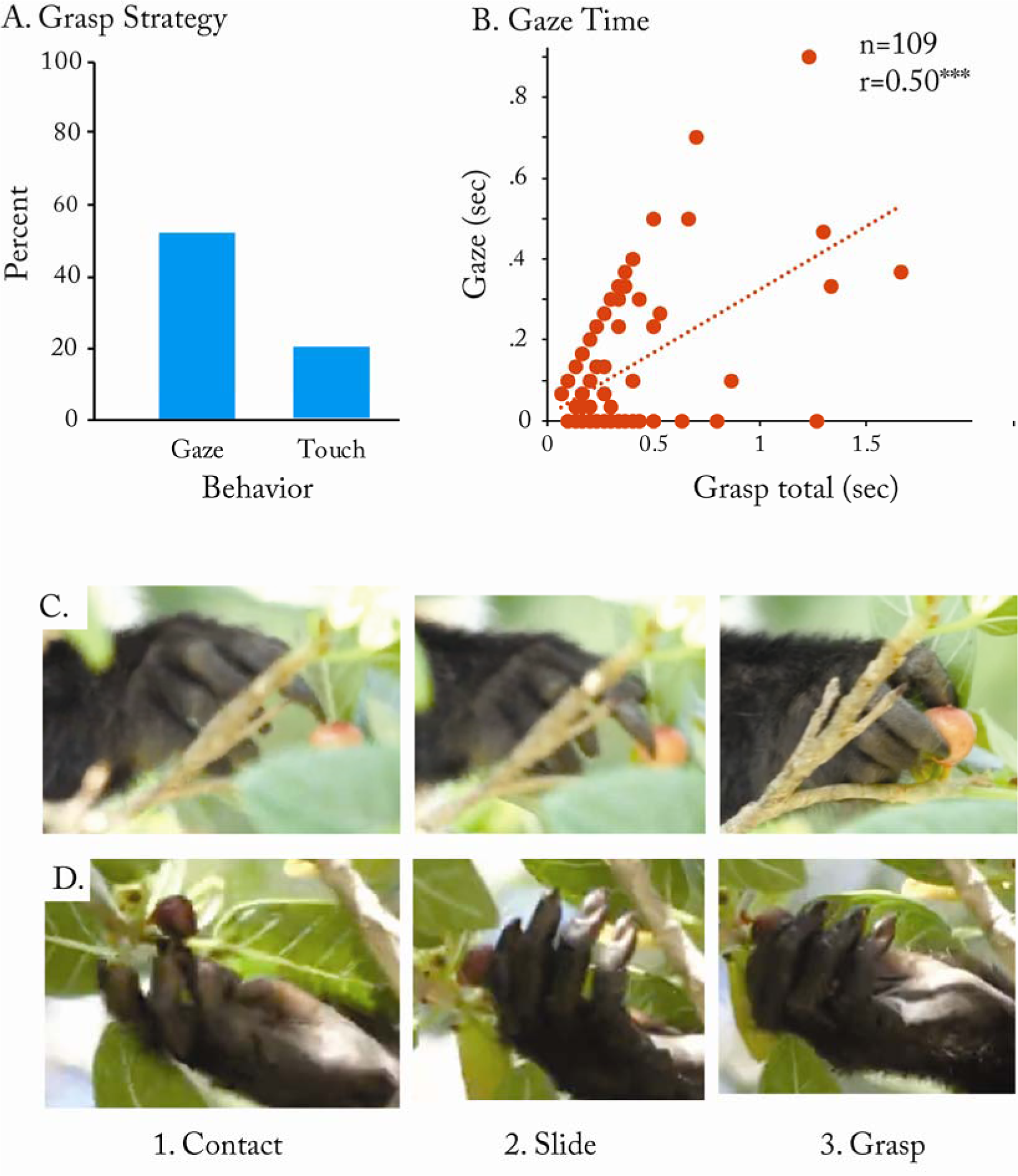
Grasp and gaze. (A). Grasp strategy shows that about half of all reaches were not associated with concurrent gaze and that about a quarter of all reaches featured hand touches to the target before grasping. (B). There is a significant relationship between total gaze time and total grasp time, but the correlation is not high as about half of reaches are not associated with concurrent gaze. (C). Example of a precision grasp made with a pronated hand in which the fruit Ficus ovalis was touched by a finger that appeared to guide the grasp. (D). An example of a grasp in which a supinated hand appears to identify a *Ficus ovalis* fruit item location using touch, which then appears to guide the grasp.

### Gaze and somatosensation associated with immediate withdraw

Figure 6A illustrates that when reaching for food, the capuchins used one hand to reach for a food item for approximately half of their reaches (n=304), and for the other half of their reaches used the assistance of a second hand (n=293). As is illustrated in Figure 6B, when only one hand was used, the other hand was used for support, either on the substrate or holding onto a branch. As is illustrated in Figure 6C, when two hand were used, one hand grasped a branch containing fruit and manipulated it into a position from which the other hand could grasp the fruit. Despite the grasping strategies used to obtain fruit items, they were always brought to the mouth with one hand.

**Figure 6.**
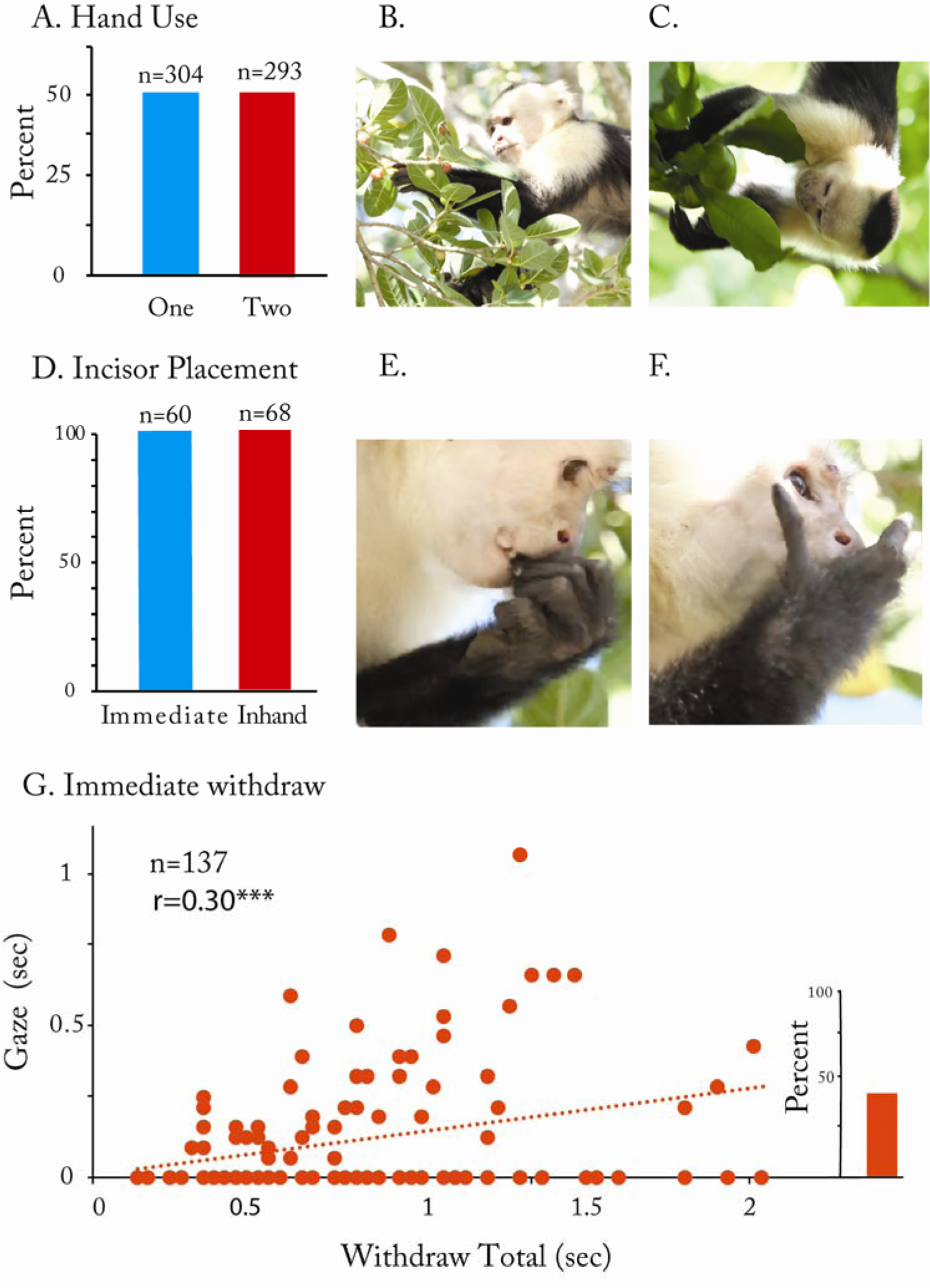
Hand use during reach and withdraw. (A). One or both hands could participate in fruit picking as only one hand picked whereas the other provided support or one manipulated a branch as the other reached. (B). A capuchin named Badger uses one hand to reach. (C) Badger uses one hand to manipulate a branch and the other hand to reach. (D). Fruit was grasped with the incisors when presented to the mouth with a precision grasp or a whole hand grap. A capuchin named Palpatine (PA) presents a fruit item to the mouth with a precision grasp (E) or a whole hand grasp (F). Note that with the precision grasp the hand was partially supinated (palm-vertical) and with a whole hand grasp the hand is fully supinated (the palm the facing mouth). G. Relation between gaze directed to the fruit during immediate withdraw and total withdraw time showing that the relation was weakly significant (p=0.30) because almost half of immediate withdraws were not associated with gaze (insert).

Figure 6D summarizes the frequency with which fruit items were placed between/grasped by the incisors upon immediate withdraw. For 60 of 128 immediate withdraws, the item was grasped with a precision grasp and transferred to the incisors with the palm in a relatively vertical orientation. Figure 6E shows the transfer of food with the hand with the item held in a precision grip with the hand in a 90^0^ supinated position. For 69 of 128 withdraws, the food was held in a whole hand grasp and was transferred to the incisors with the hand fully supinated as is shown in Figure 6F. Transfers of food items from the hand to the mouth involved one relatively discrete movement, with few transfers involving repeated bites of the food item, and no transfers missing placement to the incisors and requiring a correction. Nevertheless, most observations of food transfer involved the transfer of fruit items that were easy to take with a bite. There were instances in which the capuchins were observed to grasp items with the premolars, but these were either instances in which they were trying to dislodge a fruit item such as *Bromelia pinguin* (a food item growing in clusters) with their mouth or were biting the bark from a stick with the mouth. These activities were not formally investigated.

As is shown in Figure 6D-E, precision grasps were used to grasp food items that were small (e.g., *Ficus ovalis* and *Ficus cotinifolia*, which are grape sized) and whole hand grasps to obtain food items that are larger (e.g., *Spondias mombin*and *Diospyros salicifolia*, which are plum sized). Of 89 grasps, 42% (37/89) were made using a precision grip and 58% (49/89) were whole hand grips. A correlation between grasping posture and food size gave a significant correlation between grip type and food size, r(87) = 0.74, p<0.001. Of the 89 observations, there were 10 where precision grips were used for larger food items and one where a whole hand grip was used for a small food item.

Figure 6G shows the relationship between gaze and total time taken to grasp a food item for an immediate withdraw. Of 139 observations for which the face was oriented so gaze could be ascertained, there was a weak association between gaze duration and time to withdraw, r(137)=0.30, p<0.05. As is shown by the insert in Figure 6D, less than 50% of immediate withdraws were associated with gaze directed to the food item during the withdraw. A similar absence of an association between gaze and immediate withdraw is reported for macaques and for humans (de Bruin et al., 2008; Hirsche et al, 2022; Sacrey et al., 2011).

An examination of inhand withdraws showed that for 94% (169/179) of the withdraw movements, the food was held in a precision grip (Figure 3E). Because 97% of inhand holding movements were with larger food items, at least one objective of inhand manipulations involved moving the object from a whole hand grasp to a precision grasp. Of 104 inhand withdraws in which food was transferred to the mouth and a bite was taken from the food, 87 (83.6%) were associated with hand adjustments including releasing and regrasping the food item so that it was placed/replaced in a precision grip.

### Gaze, precision grip and hand posture associated with inhand withdraw

Figure 7 illustrates hand use for inhand holding and withdrawing of a food item to the mouth and the relation of gaze to holding and withdraw movements. The objective of withdraw actions appears to be obtaining end point comfort of the optimal placement of the food in the mouth (Rosenbaum et al, 2012; Truppa et al, 2020). From 417 inhand withdraw movements, holding was most frequently associated with two hands (Figure 7C), whereas withdrawing the food item to the mouth was associated with one hand (Figure 7D). The shift from two hands to one hand, as shown in Figure 7C, yielded a significant Pearson Chi-Square value, Chi-Square(1)=175, p<0.001. The significant effect suggests a preference for two-hand holding vs. one-hand withdraw for the fruit items that were being eaten.

**Figure 7.**
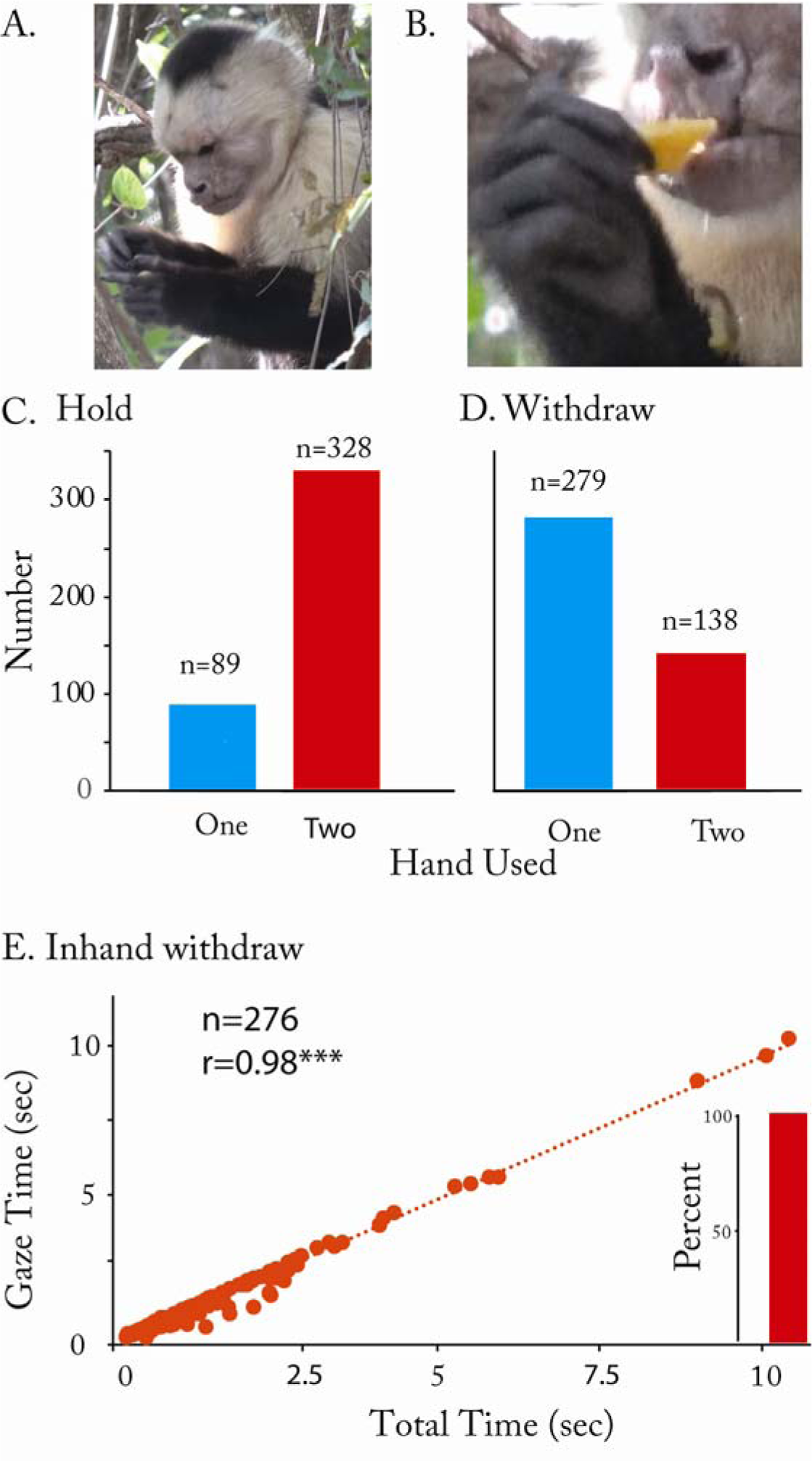
Inhand withdraw and gaze. (A). A capuchin named by Cicatriz (CZ) manipulates a fruit item to place it into a precision grasp under gaze. (B) Cicatriz (CZ) presents the food item held with a precision grasp and with a partially supinated hand (palm-vertical) to the mouth. (C). The high incident of two-handed fruit handling. (D). The high incidence of one handed withdraw. (E). The relation between gaze time and total handling and withdraw time was significant (r(273)=0.98) showing that gaze related activity accounted for most of the food handling/withdraw time. The insert (Percent) shows that all food handling events were associated with gaze directed to the food manipulation movements.

During food holding, a food item could be passed between the hands, picked at with one hand while being held with the other, or manipulated to an orientation of comfort so it could be placed in the mouth. An analysis of the grasp used while withdrawing the food to the mouth showed that for 391 of 417 (94.4%) of the withdraws, the food item was held in a precision grip and the palm was in a 90° orientation, with a palm-in orientation, for food transfer to the mouth. Because most of the food items taken from an inhand holding position were relatively large, this result suggests that the objective in handling the food item was to regrip an item so that the item was held with a precision grip for presentation to the mouth. As an assist in maintaining this grip and hand posture, it was observed that as the capuchins took a bite from a food item, they also frequently adjusted their hand position, by quickly releasing and regrasping the item, to maintain a precision grip (see above). On those instances on which both hands withdrew a food item to the mouth, a precision grip was used and both hands were similarly oriented in a palm-vertical posture.

Measures of the duration of gaze anchoring on the food item, including holding/manipulating and withdrawing relative to the total duration of holding and withdrawing, were obtained from 279 holding and withdraw events is illustrated in Figure 7E. (The number of withdraw movements associated with gaze anchoring measures was limited to those in which the face and eyes could be seen.) Because gaze was only disengaged for a brief and relatively constant duration of time during the latter portion of the withdraw, nearly all variation in time, illustrated in Figure 7E, is associated with the period of gaze anchoring. This relationship was supported by a significant correlation, r(275)=0.98, p<0.001, between gaze duration and total handling and withdraw time. Gaze was also maintained during the initial portion of the withdraw itself, and there was a significant relationship between gaze duration during withdraw and total withdraw time, r(275)=.709, p<0.001. The average point of occurrence of eye disengage during the withdraw was at 70.1% of the way through the withdraw (between the first frame of the video where the wrist began to approach the mouth to the point that wrist movement stopped as the food was transferred to the incisors).

### A blink indicated the point of eye disengage

A blink, as described in humans and macaques (de Bruin, et al, 2008; Hirsche, 2022), was reliably associated with gaze/head disengage during inhand withdraw movements as shown in Figure 8 and in Video 3. Figure 8A illustrates a blink by a capuchin that occurred just as gaze was disengaged during an inhand withdraw movement, after a capuchin had fished some pulp from a fruit with its index finger. As illustrated in Figure 8B, the blink was associated with raising the head from a downward orientation, in which gaze was directed to the food, to a horizontal position, in which the food was accepted by the mouth. Figure 8C illustrates that the relationship between immediate withdraw and blinks was not as clear as it was for inhand withdraw and blinks. Of 55 immediate withdraws in which the face was visible, only half were associated with a disengage blink, whereas of 98 inhand withdraws, 96 were all associated with a disengage blink. As noted in the section related to Figure 7, for immediate withdraws, gaze disengage often occurred before an item was grasped and so might be missed if the head was not oriented to the camera. For inhand withdraws, the blinks were more seemingly time influenced and were initiated approximately at a point 67% through the withdraw (the initiation of a blink preceded head disengage), as the hand transitioned from holding and manipulating a food item to placing the food item in the mouth.

**Figure 8.**
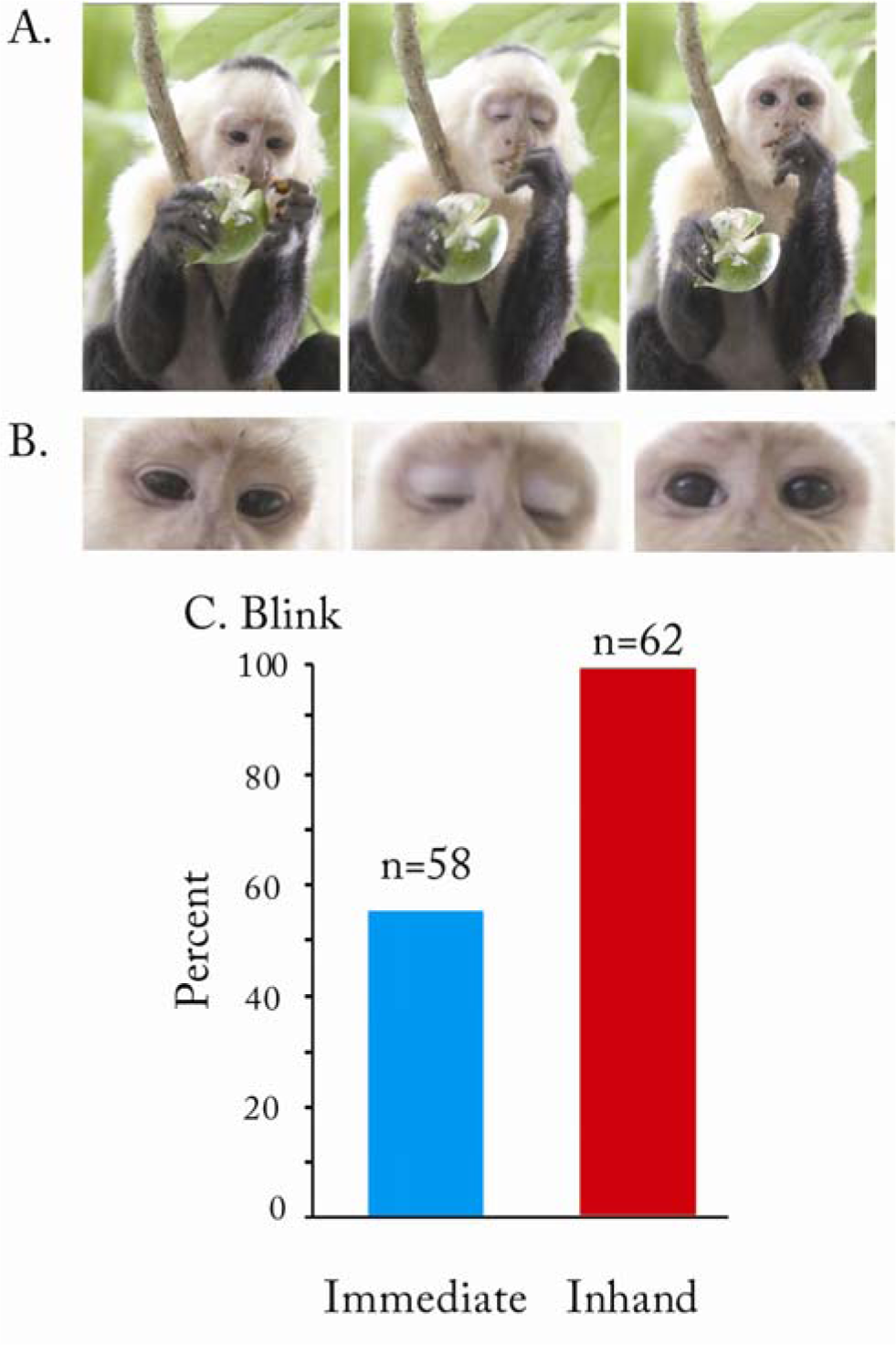
Blink and withdraw. (A). A capuchin named Appo (AP) uses a finger to fish pulp from *Stemmadenia obovata* and withdraw the pulp to the mouth with an index finger. In the sequence the finger is inserted into the middle of the fruit pulp to obtain a piece of pulp and then the finger takes the pulp to the mouth. (B). Finger fishing is associated with gaze anchoring and gaze is disengaged with an associated eye blink during the withdraw to the mouth. (C). The percent of blinks with immediate withdraw were not high as blinks could occur before a fruit item was grasped, vs. a high incidence of blinks occurred with inhand withdraw because gaze was always directed to the fruit when it was handled.

## Discussion

This study asked whether the platyrrthine primate, *Cebus imitator*, used a visually mediated strategy to orient food items in the hand for withdraw to the mouth, as do catharrhine primates, or whether it used a somatosensory strategy of reaching with the mouth to take food from the hand, as do strepsirrhine primates. While engaged in arboreal fruit foraging, the capuchins used a sitting posture for fruit picking and eating, even though foraging in the teerminal branches of trees. Bilateral hand movements, in which one hand manipulates a branch as the other hand reaches for food, contribute to fruit picking. Reaching component movements (the reach, the grasp, and the withdraw) are used flexibly with respect to spatial, temporal, and interactive features, suggesting independence in their control. The capuchins used two types of withdraws, an immediate withdraw, bringing food directly to the mouth after it is grasped, and an inhand withdraw, positioning food in the hand for presentation to the mouth. Vision plays a central role in the food handling of inhand withdraw.

Immediate withdraw movements were sometimes associated with gaze directed to the target food item but just as frequently occurred without gaze maintained on the food item. Although the capuchins might glance at an item before reaching, disengage often occurred before a food item was grasped and the grasp itself was often associated with target touching (see Melin et al, 2022 for a description touch use in by capuchin monkeys). Nevertheless, small items were grasped with a precision grip and large items with a whole hand grip, and items of both sizes could be brought to the mouth with an immediate withdraw without visual monitoring. That food items can be grasped and brought to the mouth without visual monitoring has been reported for humans fitted with eyetracking glasses. The same studies show that visual occlusion does not disrupt the accuracy of this form of withdraw, supporting the idea that it is mediated largely by somatosensation (de Bruin et al., 2008; Karl et al., 2012; Sacrey et al., 2011).

Food items held inhand were associated with visual monitoring, whereby they were manipulated until they were held in a precision grip for withdraw to the mouth. As the hand was brought to the mouth, it was oriented to a palm-vertical posture that exposed the food item protruding from the radial aspect of the fingers to be grasped by the incisors. Even though gaze was maintained during inhand manipulation, it was disengaged partway through the withdraw. The disengage was associated with a blink and raising of the head so that the mouth was horizontal when it took the food with a bite by the incisors. Consequently, although food orienting and the initial phase of the withdraw were monitored with vision, placement in the mouth was mediated by somatosensation as occurred for an immediate withdraw. If a large food item was brought to the incisors, the hand could be repositioned as the incisors held the food, presumably to improve the precision grip, an action that would assist in positioning the food item for the next withdraw. Despite repositioning the food item in the hand as it is presented to the mouth, subsequent withdraw movements were still associated with gaze/disengage behavior indicating that the location and trajectory of the food to the mouth is visually confirmed on every withdraw. Just as catarrhine primates use vision and a variety of hand orientations to present food to the mouth (Hirsche et al, 2022), capuchins also use vision but also rely on a precision grasp with a vertical hand orientation, suggesting a simpler form of withdraw.

It has been suggested that the visual control of hand shaping initially evolved for insect or fruit harvesting in stem primates occupying a fine-branch niche (Cartmill, 1972, 2012; Sussman & Raven, 1978; Sussman et al., 2013; Scott, 2019). Because strepsirrhine primates do not use visual mediated inhand food manipulation, whereas platyrrhine primates (as shown here) and catarrhine primates (Hirsche et al, 2022) do, it is likely that a visually mediated withdraw evolved in a stem lineage preceding platyrrhine-catarrhine divergence. Possibly the visual manipulation of objects held in the hand was then adopted for visually mediated object grasping in platyrrhines and catarrhines (Hirsche et al, 2022). The objective of inhand food manipulation and precision grasping is likely the attainment of end point comfort; i.e., the optimal placement of food in the mouth (see Rosenbaum et al, 2012 for a review of the relationship between the way that an object is grasped and endpoint comfort). Although capuchins do not display the variety of hand shaping movements of catarrhine primates (see, Spinozzi et al, 2004; Truppa, et al, 2019; 2021 for descriptions of capuchin grasping), the visual control of precision grasping likely led to the many modifications documented in platyrrhines and catarrhines. Taken together, it is evident that the visual control of objects using the hands evolved in stages in which the reach, grasp and withdraw have their own evolutionary histories as is suggeted by visuomotor channel theory (Jeannerod,1981; Jeannerod et al., 1995, 1998; Grant & Conway, 2019; Sartori et al, 2015; Whishaw et al, 2016).

The association of eye-disengage with an eye blink has also been described in humans and macaques (de Bruin et al., 2008; Hirsch et al, 2022; Karl et al., 2012; Sacrey et al., 2011). Eye blinking occurred in capuchins and its onset is closely associated with the head disengage movement. Proposed explanations for blinks include the ideas that they are associated with visual focus and/or related shifts in brain networks. With respect to focus, the idea is that visualizing a target held in the hands requires accommodation (Kiorpes, 2019), and a blink may represent a release from the strain of accommodation (Ang and Maus, 2020) and/or may facilitate focusing when looking elsewhere (Jaschinski et al., 1996). Blinks have been associated with a brain network change from an active to a default mode (Brych & Händel, 2020; Nakano et al., 2013). In relation to this idea, a blink may assist in removing visual attention from a target that is approaching the mouth to somatosensory control by which it is placed in the mouth. Because strepsirrhines do not display a withdraw-associated blink, blink-associated disengage likely also appeared before the catarrhine-platyrrhine divergence in association with visual inspection and manipulation of food items held in the hand.

A caveat of the visual object monitoring of *Cebus imitator*, as described here, relates to the similarities of many of the food items for which they were foraging. The figs and plum-sized fruits although varying in size, were round and so might not be representative of food items that protrude from the hands. Incidental observations of the handling of other items did show that the capuchins could use other handling strategies. When holding sticks upon which they chewed, the capuchins usually brought their mouth to the stick rather than bringing the stick to the mouth. They also brought their mouth to grasp *Luehea candida*, a large shell containing seeds, when attempting to lick seeds from them. When shaking *Luehea candida* shells to remove their seeds, capuchins were also observed to bring freed seeds to the mouth in an open palm, and when removing aril from *Stemmadenia obovata*, they used only an index finger. Nevertheless, for these behaviors including shaking objects and using a single digit, they usually used a palm-vertical strategy similar to that used when withdrawing fruit to the mouth. Future studies could further investigate whether eating/handling of a wider range of objects is associated with additional hand postures (Truppa et al, 2021).

In conclusion, the question that motivated the present research was whether platyrrhine anthropoids use visual strategies to assist their withdraw of food items, as do macaques and other catarrhine anthropoids (Hirsche et al, 2022). The present results show that the visual manipulation of food items during withdraw indeed occurs in platyrrhines. Like catarrhines, capuchins gaze at food items as they orient them for the withdraw, they disengage vision partway through the withdraw with a blink, and they raise the mouth to horizontal to take the food with the incisors. Unlike catarrhines, capuchin precision grips are simpler, and they rely on a palm vertical withdraw. Our video repositories of other platyrrhine primates suggest that this behavior is not unique to capuchins and so likely originates prior to the platyrrhine-catarrhine divergence and so predates the precision grasps used by catarrhines. Amongst the studies on the neural control of hand use, the finding that there is a deep penetration of the direct projection of the corticospinal tract to cervical motor neurons in capuchins, as also occurs in catarrhines, has been attributed to their use of finger shaping movements when grasping (Bortoff and Strick, 1993). The present results raise the possibility that these projections may be additionally related to visually mediated inhand food positioning. The association between inhand and grasp-related object manipulation and their neural control could be further investigated in comparative studies of the many other platyrrhine monkey species.

## Acknowledgements

We thank R. Blanco Segura and M.M. Chavarria and staff from the Área de Conservación Guanacaste and Ministerio de Ambiente y Energía. Warmest thanks also to Saúl Cheves Hernandez and Ronald Lopez Navarro for field assistance. Funding was generously provided by the National Science Foundation’s Biological Anthropology program (BCS-1945771, BCS-1944915, BCS-1945283, BCS-1945767). This research adhered to the laws of Costa Rica, the United States, and Canada and complied with protocols approved by the Área de Conservación Guanacaste and by the Canada Research Council for Animal Care through the University of Calgary’s Life and Environmental Care Committee and by Mercer University’s Institutional Animal Care and Use Committee.

## Videos

**Video 1**. A reach in which a capuchin named Badger is eating *Ficus ovalis*. The capuchin visually disengages a fruit item with a blink and head movement that brings the mouth to a vertical orientation before completing the reach and making, a precision grasp and a withdraw. Note the mouth gape as the hand advances to the fruit target and the spitting movements during the withdraw movement. Note also that the right manipulates the branch so that the fruit can be reached for with the left hand.

**Video 2**. A capuchin named MonMothma (MT) makes an underhand touch-associated precision grasp. The target fruit *Ficus ovalis* is first touched and then the hand reverses movement direction in order to make a precision grasp of the item.

**Video 3**. A capuchin named Appo (AP) pulp fishes with a finger in *Stemmadenia obovata* fruit. Appo makes a withdraw associated with visual disengage, a blink and a lift of the mouth to a horizontal orientation of accept the food item from the finger with the incisors.

## References

Ang, J.W.A., & Maus, G.W. (2020). Boosted visual performance after eye blinks. Journal of Vision, 1, 20–22. https://doi.org/10.1167/jov.20.10.2

Arnold C, Matthews LJ, Nunn CL. The 10kTrees website: a new online resource for primate phylogeny. Evolutionary Anthropology: Issues, News, and Reviews. 2010 May;19(3):114–8.

Bishop, A. (1964). Use of the hand in lower primates. In J. Buettner-Janisch (Ed.). Evolutionary and genetic biology of primates (pp. 135–225). Academic Press.

Bortoff GA, Strick PL. Corticospinal terminations in two new-world primates: further evidence that corticomotoneuronal connections provide part of the neural substrate for manual dexterity. J Neurosci. 1993 Dec;13(12):5105–18. doi: 10.1523/JNEUROSCI.13-12-05105.1993. PMID: 7504721; PMCID: PMC6576412

Brych, M., & Händel, B. (2020). Disentangling top-down and bottom-up influences on blinks in the visual and auditory domain. International Journal of Psychophysiology, 158, 400–410. https://doi:10.1016/j.ijpsycho.2020.11.002

Cartmill, M. (1972). Arboreal adaptations and the origin of primates. In R. Tuttle (Ed.), The functional and evolutionary biology of primates (pp. 97–122). Aldine-Atherton.

Cartmill, M. (1974). Rethinking primate origins. Science, 184, 436e443. https://doi:10.1126/science.184.4135.436

Cartmill, M. (1992). New views on primate origins. Evolutionary Anthropology, 1, 105e111. doi:10.1002/evan.1360010308

Cartmill, M. (2012). Primate origins, human origins, and the end of higher taxa. Evolutionary Anthropology, 21, 208e220. https://doi.org/10.1002/evan.21324

Christel, M. (1993). Grasping techniques and hand preferences in *Hominoidea*. In H. Preuschoft & D. J. Chivers (Eds.), Hands of primates (pp. 91–108). Springer.

Christel, M., & Fragaszy, D. (2000). Manual function in *Cebus apella*. Digital mobility, preshaping, and endurance in repetitive grasping. International Journal of Primatology, 21, 697–719. https://doi.org/10.1023/A:1005521522418

de Bruin, N., Sacrey, L.A., Brown, L.A., Doan, J., & Whishaw, I.Q. (2008). Visual guidance for hand advance but not hand withdrawal in a reach-to-eat task in adult humans: reaching is a composite movement. Journal of Motor Behavior, 337346. https://doi.org/10.3200/JMBR.40.4.337-346

Edwards, M.G., Wing, A.M., Stevens, J., & Humphreys, G.W. (2005). Knowing your nose better than your thumb: measures of over-grasp reveal that face-parts are special for grasping. Experimental Brain Research, 161, 72–80. https://doi:10.1007/s00221-004-2047-2

Fragaszy, D.M. (1968). How non-human primates use their hands. In K. Connolly (Ed.), Psychobiology of the hand (pp. 77–96). MacKeith Press.

Grant. S., & Conway, M.L. (2019) Some binocular advantages for planning reach, but not grasp, components of prehension. Experimental Brain Research, 237, 1239–1255. https://doi.org/10.1007/s00221-019-05503-4

Hallgren, K.A. (2012). Computing inter-rater reliability for observational data: an overview and tutorial. Tutorials in Quantative Methods in Psychology, 6, 23–34. doi: 10.20982/tqmp.08.1.p023

Hirsche, L. A., Cenni, C., Leca, J.-B., & Whishaw, I. Q. (2022). Two types of withdraw-to-eat movement related to food size in long-tailed macaques (Macaca fascicularis): Insights into the evolution of the visual control of hand shaping in anthropoid primates. Animal Behavior and Cognition, 9(2), 176–195. https://doi.org/10.26451/abc.09.02.02.2022

Iwaniuk, A.N., Ivanco, T.L., Nelson, J.E., Pellis, S.M., & Whishaw, I.Q. (1998) Reaching, grasping and manipulation of food objects by two species of tree kangaroos, *Dendrolagus lumholtzi* and *Dendrolagus matschiei*. Australian Journal of Zoology, 46, 235–248. https://doi:10.1071/ZO98004

Iwaniuk, A.N., & Whishaw, I.Q. (2000). On the origin of skilled forelimb movements. Trends in Neuroscience, 23, 372–376. https://doi:10.1016/s0166-2236(00)01618-0

Ivanco, T.L., Pellis, S.M., Whishaw, I.Q. (1996). Skilled forelimb movements in prey catching and in reaching by rats (*Rattus norvegicus*) and opossums (*Monodelphis domestica*): Relations to anatomical differences in motor systems. Behavioural Brain Research, 79, 163–81. https://doi:10.1016/s0166-2236(00)01618-0

Jeannerod, M, (1981). Intersegmental coordination during reaching at natural visual objects. In J. Long & A. Badeley (Eds.), Attention and performance IX (pp. 153–169). Lawrence Erlbaum Associates.

Jeannerod, M., Arbib, M.A., Rizzolatti, G., & Sakata, H. (1995). Grasping objects: the cortical mechanisms of visuomotor transformation. Trends in Neuroscience, 18, 314–20.

Jeannerod, M., Paulignan, Y., Weiss, P. (1998). Grasping an object: One movement, several components. Novartis Foundation Symposium, 218, 5–16.

Karl, J.M., & Whishaw, I.Q. (2013). Different evolutionary origins for the reach and the grasp: an explanation for dual visuomotor channels in primate parietofrontal cortex. Frontiers in Neurology, 234, 208. https://doi.org/10.3389/fneur.2013.00208

Karl, J.M., Sacrey, L.A., Doan, J.B., & Whishaw, I.Q. (2012) Oral hapsis guides accurate hand preshaping for grasping food targets in the mouth. Experimental Brain Research, 221, 223–40. https://doi.org/10.1007/s00221-012-3164-y

Kay, R.F., Ross, C., & Williams, B.A. (1997). Anthropoid origins. Science, 275, 797–804. https://doi:10.1126/science.275.5301.797

Kelley, K., & Preacher, K. J. (2012). On effect size. Psychological Methods, 17, 137–152. https://doi.org/10.1037/a0028086

Kiorpes, L. (2019). Understanding the development of amblyopia using macaque monkey models. Proceedings of the National Academy of Science, 116, 26217–26223. https://doi:10.1073/pnas.1902285116.

Laird, M.F., Punjani, Z., Oshay, R.R., Wright, B.W., Fogaça, M.D., van Casteren, A., Izar, P., Visalberghi, E., Fragazy, D., Strait, D.S. and Ross, C.F. (2022). Feeding postural behaviors and food geometric and material properties in bearded capuchin monkeys (*Sapajus libidinosus*). American Journal of Biological Anthropology, 178(1), pp.3–16. https://doi.org/10.1002/ajpa.24501

Macfarlane, N.B., & Graziano, M.S. (2009). Diversity of grip in *Macaca mulatta*. Experimental Brain Research, 197, 255–68. https://doi:10.1007/s00221-009-1909-z

Marzke, M.W., Marchant, L.F., McGrew, W.C., Reece, S.P. (2015). Grips and hand movements of chimpanzees during feeding in Mahale Mountains National Park, Tanzania. American Journal of Physical Anthropology, 156, 317–326. https://doi.org/10.1002/ajpa.22651

Melin AD, Veilleux CC, Janiak MC, Hiramatsu C, Sánchez-Solano KG, Lundeen IK, Webb SE, Williamson RE, Mah MA, Murillo-Chacon E, Schaffner CM, Hernández-Salazar L, Aureli F, Nakano, T., Kato, M., Morito, Y., Itoi, S., & Kitazawa, S. (2013). Blink-related momentary activation of the default mode network while viewing videos. Proceedings of the National Academy of Science, 110, 702–706. https://doi.org/10.1073/pnas.1214804110

Peckre, L.R., Fabre, A.-C., Hambuckers, J., Wall, C.E., Socias-Martînez, L., & Pouydebat, E. (2019). Food properties influence grasping strategies in strepsirrhines. Biological Journal of the Linneal Society, 20, 1–55.

Peckre, L.R., Fabre, AC., Wall, C.E. et al. Evolutionary History of food Withdraw Movements in Primates: Food Withdraw is Mediated by Nonvisual Strategies in 22 Species of Strepsirrhines. Evol Biol (2023). https://doi.org/10.1007/s11692-023-09598-0

Pessina MA, Bowley BGE, Rosene DL, Moore TL. A method for assessing recovery of fine motor function of the hand in a rhesus monkey model of cortical injury: an adaptation of the Fugl-Meyer Scale and Eshkol-Wachman Movement Notation. Somatosens Mot Res. 2019 Mar;36(1):69–77. doi: 10.1080/08990220.2019.1594751. PMID: 31072219; PMCID: PMC6643292.

Posner, M.I., Inhoff, A.W., Friedrich, F.J., & Cohen A. (1987). Isolating attentional systems: A cognitive-anatomical analysis. Psychobiology, 15, 107–121. https://doi.org/10.3758/BF03333099

Pouydebat, E., Laurin, M., Gorce, P., & Bels, V. (2008). Evolution of grasping among anthropoids. Journal of Evolutionary Biology, 21, 1732–1743. https://doi.org/10.1111/j.1420-9101.2008.01582

Pouydebat, E., Gorce, P., Coppens, Y., & Bels, V. (2009). Biomechanical study of grasping according to the volume of the object: Human versus non-human primates. Journal of Biomechanics, 42, 266-72. https://doi:10.1016/j.jbiomech.2008.10.026.

Reghem, E., Tia, B., Bels, V., Pouydebat, E. (2011). Food prehension and manipulation in *Microcebus murinus* (*Prosimii, Cheirogaleidae*). Folia Primatologica (Basel), 82, 177–88. doi: 10.1159/000334077.

Rosenbaum DA, Chapman KM, Weigelt M, Weiss DJ, van der Wel R. Cognition, action, and object manipulation. Psychol Bull. 2012 Sep;138(5):924-46. doi: 10.1037/a0027839. Epub 2012 Mar 26. PMID: 22448912; PMCID: PMC3389205.

Sacrey, L. R., Karl, J. M., & Whishaw, I. Q. (2012). Development of rotational movements, hand shaping, and accuracy in advance and withdrawal for the reach-to-eat movement in human infants aged 6-12 months. Infant Behavioral Development, 35, 543–560. https://doi:10.1016/j.infbeh.2012.05.006

Sacrey, L.A., Travis, S.G., & Whishaw, I.Q. (2011). Drug treatment and familiar music aids an attention shift from vision to somatosensation in Parkinson’s disease on the reach-to-eat task. Behavioural Brain Research, 217, 391-388. https://doi:10.1016/j.bbr.2010.11.010.

Sartori, L., Camperio-Ciani, A., Bulgheroni, M., Castiello, U. (2015). Intersegmental coordination in the kinematics of prehension movements of macaques. PLoS One, 10(7):e0132937. doi: 10.1371/journal.pone.0132937. PMID: 26176232; PMCID: PMC4503540.

Scott, J.E. (2019). Macroevolutionary effects on primate trophic evolution and their implications for reconstructing primate origins. Journal of Human Evolution, 133, 1–12. https://doi:10.1016/j.jhevol.2019.05.001.

Spinozzi G, Truppa V, Laganà T. Grasping behavior in tufted capuchin monkeys (Cebus apella): grip types and manual laterality for picking up a small food item. Am J Phys Anthropol. 2004 Sep;125(1):30–41. doi: 10.1002/ajpa.10362. PMID: 15293329.

Sussman, R.W (1991). Primate origins and the evolution of angiosperms. American Journal of Primatology, 23, 209e223. https://doi.org/10.1002/ajp.1350230402

Sussman, R.W., Rasmussen, D.T., & Raven, P.H. (2013). Rethinking primate origins again. American Journal of Primatology, 75, 95e106. https://doi.org/10.1002/ajp.22096

Sussman, R.W., & Raven, P.H. (1978). Pollination by lemurs and marsupials: An archaic coevolutionary system. Science, 200, 731e736. https://doi:10.1126/science.200.4343.731.

Sustaita, D., Pouydebat, E., Manzano, A., Abdala, V., Herrel, F., & Herrel, A. (2013). Getting a grip on tetrapod grasping: Form, function, and evolution. Cambridge Review of the Cambridge Philosophical Society, 88, 380–405. https://doi.org/10.1111/brv.12010

Truppa V, Paola Carducci, Gloria Sabbatini, (2019). Object grasping and manipulation in capuchin monkeys (genera *Cebus* and *Sapajus*), Biological Journal of the Linnean Society, 127, 563–582, https://doi.org/10.1093/biolinnean/bly131

Truppa V, Sabbatini G, Izar P, Fragaszy DM, Visalberghi E, (2021). Anticipating future actions: Motor planning improves with age in wild bearded capuchin monkeys (Sapajus libidinosus). Developmental Sciences. 24,e13077. doi: 10.1111/desc.13077.

Vainio L. (2019). Connection between movements of mouth and hand: Perspectives on development and evolution of speech. Neurosciences and Biobehavioral Reviews, 100, 211–223. doi: 10.1016/j.neubiorev.2019.03.005.

Whishaw, I.Q. & Coles, B. (1996) Varieties of paw and digit movement during spontaneous food handling in rats: Postures, bimanual coordination, preferences, and effect of forelimb cortex lesions. Behavioural Brain Research, 77, 135–148. https://doi:10.1016/0166-4328(95)00209-x

Whishaw, I.Q., Faraji, J., Mirza Agha, B., Kuntz, J.R., Metz, G.A.S., & Mohajerani, M.H. (2018). A mouse’s spontaneous eating repertoire aids performance on laboratory skilled reaching tasks: A motoric example of instinctual drift with an ethological description of the withdraw movements in freely-moving and head-fixed mice. Behavioural Brain Research, 337, 80–90. https://doi:10.1016/j.bbr.2017.09.044

Whishaw, I.Q., Ghasroddashti, A., Mirza Agha, B., & Mohajerani, M.H. (2020). The temporal choreography of the yo-yo movement of getting spaghetti into the mouth by the head-fixed mouse. Behavioural Brain Research, 381, 112241. https://doi:10.1016/j.bbr.2019.112241

Whishaw, I.Q., & Karl, J.M. (2014). The contribution of the reach and the grasp to shaping brain and behaviour. Canadian Journal of Experimental Psychology, 68, 223–235. https://doi.org/10.1037/cep0000042

Whishaw, I.Q., & Karl, J.M. (2019). The evolution of the hand as a tool in feeding behavior: the multiple motor channel theory of reaching. In V. Bels & I.Q. Whishaw (Eds.), Feeding in Vertebrates (pp. 159–188). Springer.

Whishaw IQ, Karl JM, Humphrey NK (2016). Dissociation of the Reach and the Grasp in the destriate (V1) monkey Helen: a new anatomy for the dual visuomotor channel theory of reaching. Experimental Brain Research, 234, 2351–2362. https://doi:10.1007/s00221-016-4640-6.

Whishaw, I.Q., Sarna, J.R., & Pellis, S.M. (1998). Evidence for rodent-common and species-typical limb and digit use in eating, derived from a comparative analysis of ten rodent species. Behavioural Brain Research, 96, 79–91. https://doi:10.1016/s0166-4328(97)00200-3

Whishaw, I.Q., Koples, J., Hirsche, L.A., & Karl, J.M. Spontaneous eating is children and adults, unpublished data.

